# Quantifying Molecular Forces with Serially Connected Force Sensors

**DOI:** 10.1101/405761

**Authors:** Y. Murad, I. T.S. Li

## Abstract

To understand the mechanical forces involved in cell adhesion, molecular force sensors have been developed to study tension through adhesion proteins. Recently, a class of molecular force sensors called tension gauge tether (TGT) have been developed that rely on irreversible force-dependent dissociation of DNA duplex to study cell adhesion forces. While the TGT offer high signal-to-noise ratio and is ideal for studying fast / single molecular adhesion processes, quantitative interpretation of experimental results has been challenging. Here we used computational approach to investigate how TGT fluorescence readout can be quantitatively interpreted. In particular we studied force sensors made of a single TGT, multiplexed single TGTs, and two TGTs connected in series. Our results showed that fluorescence readout using a single TGT can result from drastically different combinations of force history and adhesion event density that span orders of magnitude. In addition, the apparent behaviour of the TGT is influenced by the tethered receptor-ligand, making it necessary to calibrate the TGT with every new receptor-ligand. To solve this problem, we proposed a system of two serially connected TGTs. Our result shows that not only is the ratiometric readout of serial TGT independent of the choice of receptor-ligand, it is able to reconstruct force history with sub-pN force resolution. This is also not possible by simply multiplexing different types of TGTs together. Lastly, we systematically investigated how sequence composition of the two serially connected TGTs can be tuned to achieve different dynamic range. This computational study demonstrated how serially connected irreversible molecular dissociation processes can accurately quantify molecular force, and laid the foundation for subsequent experimental studies.

## Introduction

Cell adhesion is a complex process involved in many biological systems, from cell proliferation and differentiation to cell-cell communication, organism development, and disease processes. At the molecular level, cell adhesion to its environment or other cells is achieved through adhesion bonds formed by specific receptor-ligand interactions. Mechanical forces across these adhesion bonds are dynamically regulated in order for cells to sense their environment and control their functions. Therefore, monitoring the dynamics of molecular force is crucial to understanding cell adhesion.

One approach to understand molecular adhesion bond is by single molecule force spectroscopy (SMFS) where the interactions between a single pair of receptor and ligand are physically ruptured by force. This has been done through a number of high resolution techniques (atomic force microscopy, optical tweezers, and magnetic tweezers) and massively parallel techniques (centrifugal force microscopy, and acoustic force spectroscopy) (1–3). When a single adhesion bond is subjected to a constant force or increasing force at a constant force loading rate, its behaviour is characterized by either its lifetime (*τ*) or the mean rupture force (*F*_*r*_). Because bond rupturing is an intrinsically non-equilibrium process, *τ* or *F*_*r*_ measurements a single force or loading rate are insufficient to describe the full behaviour of an adhesion bond. In addition, without measuring molecular adhesion forces in living systems, it is impossible to determine the biologically relevant range of force or loading rate for the adhesion bond of interest.

The development of molecular force sensors aims to address this question. There are two major classes of force sensors using either reversible or irreversible molecular force sensing elements. Reversible force sensors can be further divided into two subcategories: analog and digital. The analog design utilizes the elastomeric response of individual polymers or proteins to measure the instantaneous forces on receptors (4–8). (Figure 1a) At the single-molecule level, they are excellent force reporters. However, in live cell situations where individual sensors cannot be spatially resolved, the analog nature of the sensors also makes it difficult to tease out heterogeneity of molecular force and adhesion bond density. In addition, the tunability of their dynamic range is limited with only a handful of proteins and polymers (5, 6, 9). Reversible digital force sensors utilize the reversible two-state structural transitions of biomolecules such as DNA-hairpin (10–12) and FL peptide (13). (Figure 1b) In these designs, the folded domain undergoes rapid structural transition between folded and unfolded state to reach a dynamic equilibrium. As force increases, the folding rate decreases while unfolding rate increases, resulting in the biomolecule spending more time in the unfolded state. The time-averaged fluorescence signal from the system is essentially digital around a single threshold force if the transition is sharp. The advantage of such design is that the fluorescence signal infers about the number of sensors above the force threshold without having to spatially resolve individual sensors. However, the information trade-off of force resolution for sensor density makes it difficult to probe the magnitude of molecular force and receptor distribution even with multiplexing such sensors (13, 14). Non-reversible force sensors such as the tension gauge tether (TGT) and the nano-yoyo utilize the force-dependent rupture of a calibrated bond as the sensor to determine the relative strength of the target receptor-ligand interaction (15–18). (Figure 1c) Fluorescence signal is “on” when the sensor is ruptured or “off” if it is intact. The digital behaviour allows one to determine the density of sensors in the ruptured state via fluorescence and create adhesion footprints that maps exactly where the adhesion events happened (17). The technique provides excellent fluorescence signal-to-noise ratio, as background signal from intact sensors can be essentially suppressed. However, quantitative interpretation of molecular force using irreversible force sensors has been challenging.

**Figure 1.**
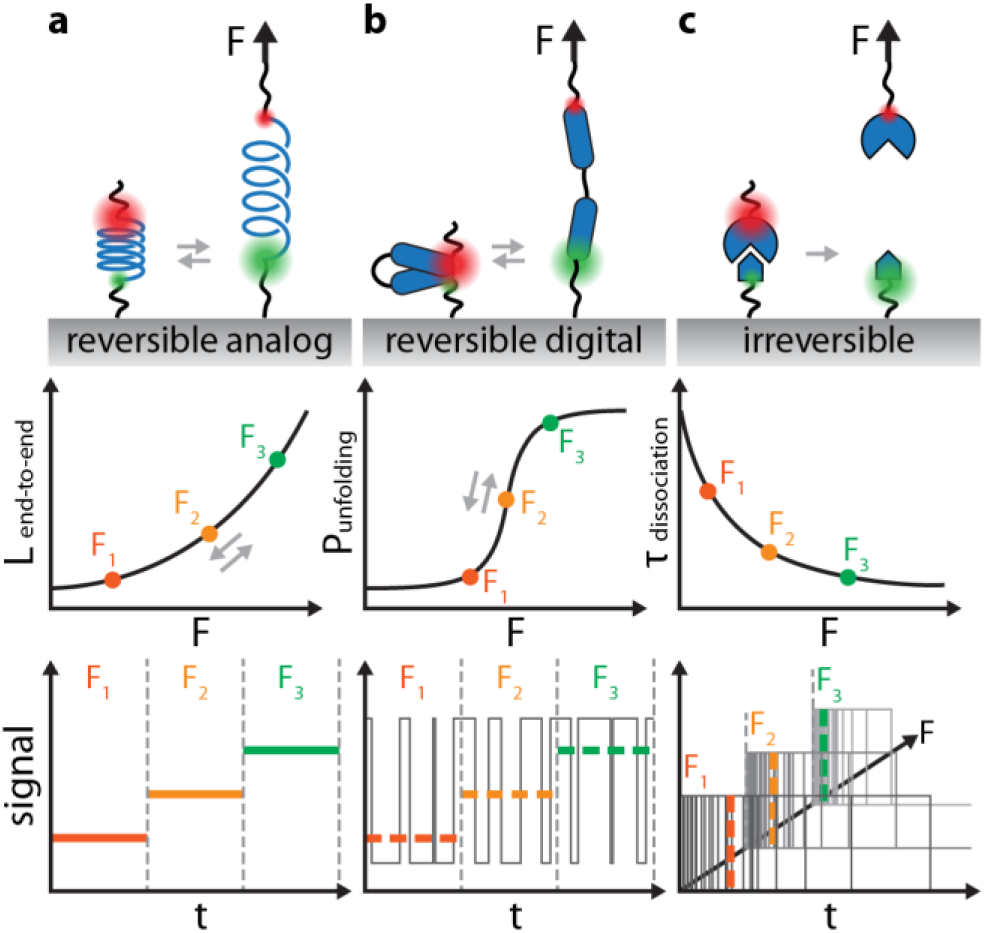
Illustration of three major classes of molecular force sensors and their fluorescence signal interpretation. **(a)** Reversible analog force sensor with an elastomer as the force sensing element (top). Elastomer end-to-end distance increases in response to tension (middle). A pair of fluorophores can produce FRET signal that directly correlate to the applied force (bottom). **(b)** Reversible digital force sensor utilizing the force-dependent two-state structural transition (such as unfolding) of a biomolecule as the force sensing element (top). The force sensing element is best described by an unfolding probability as a function of force (middle); and gives two distinct fluorescence signal levels at the single-molecule level. The dashed lines indicate the time-averaged signal level as a function of force (bottom). **(c)** Irreversible force sensors rely on the rupture of a tethered adhesion bond to inform about the relative strength of the adhesion bond of interest (top). The response of the sensor is characterized by its force dependent dissociation lifetime (middle), which can be measured from the time-dependent population statistics of the rupture events (bottom).

In this report, using simulation and TGT as an example, we first illustrate the limitations of irreversible force sensors - how the same fluorescence readout can result from drastically different mechanical histories; and how the target receptor modulates TGT characterization. We then developed a strategy using two serially connected TGT as internal calibration pairs to bypass these limitations in order to quantitatively extract force history. Lastly, we described how to design the serial TGT system is able to achieve a desirable force dynamic range.

## Methods

### Monte Carlo Simulations

Monte Carlo method was used to simulate the stochastic bond breaking of components (TGT and RGD-integrin α_5_β_1_) in the TGT-receptor complex. The simulation breaks time into short time steps (*Δt*) and assumes force is constant within each time step, while force can be arbitrarily changed between force steps to achieve any desirable force history. Since the survival probability *P* of a bond is defined by:

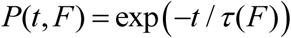

where *t* is time and *τ(F)* is the force-dependent lifetime of the bond. Force within each simulation time step is therefore, *P(Δt, F)*, and is compared to a random number between 0 and 1 to determine whether the bond survives within the time step. The simulation stops when either one of the component (TGT or RGD-integrin) breaks. The simulation code was written in MATLAB (R2017a) and ran on desktop grade CPU (Intel i7-6820HK) and GPU (NVidia GTX 980M with 1536 CUDA cores).

### Energy landscape parameters and bond lifetime calculations

We developed an energy landscape model similar to the one by Woodside et al. (19), where both the hybridization free energy and the ssDNA elastic free energy along the unzipping reaction coordinate are incorporated. In order to generalize the energy landscape parameter for a particular sequence length and CG content, the energy landscape of 100 random DNA sequences were calculated for each sequence length and CG content combination. The average unzipping energy barrier height and width were used for subsequent calculations. The energy landscapes parameters calculated this way fit closely to to the experimental DNA unzipping experimental data from Woodside et al. (19) The use of random sequences was to generalize the parameters to a given DNA sequence length and CG content.

The bond lifetime for TGT unzipping is calculated using the DNA unzipping model from Woodside et al. (19):

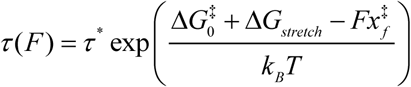

where *τ*^*\**^ *= 4.12*×*10*^*-^5^*^ *s* is the intrinsic time constant for barrier crossing; *k*_*B*_ is the Boltzmann constant; *T* is the temperature; *F* is the force applied to the bond;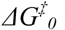is the height of the unfolding energy barrier at zero force calculated by Mfold (20); 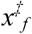is the distance to the transition state from the folded state; *ΔG*_*stretch*_ is the elastic contribution to the free energy from ssDNA stretching:

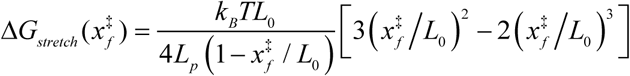

where *L*_*p*_ is the persistence length of ssDNA at 1 nm/nucleotide, *L*_*c*_ is the contour length of freed ssDNA after unzipping with 0.59 nm/nucleotideand *F*_*max*_ is the force causing the flattening of the energy landscape defined as 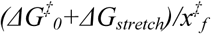. Consider N bonds are serially connected and assume no two bonds in the series rupture simultaneously, the probability that the i^th^ bond ruptures at a given force is:

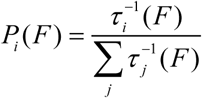

### Simulation parameters

All simulations were run at *T = 298 K*. Constant force simulations were carried at 0.1 pN increments starting from 0 pN to *F*_*max*_ for dsDNA. The temporal resolution (*Δt*) was independently determined for each force as a function of *τ(F)* as being equal to the 1/100 of the smallest *τ(F)*. Simulations were run for 10^4^ trials per simulation condition. Both types of simulation were run on the GPU when possible. All the parameters above apply unless otherwise noted.

## Results

### Monte Carlo simulation of force-dependent bond rupture kinetics

In order to simulate bond rupture behaviour, we first developed a Monte Carlo simulation to capture the stochastic rupture kinetics and statistics of a system of bonds with arbitrary force history and temporal resolution. With GPU computing and scalable time steps, we are able to probe the force-dependent dissociation behaviour of 10^5^ bonds over 12 orders of magnitude in time on a single PC. Experimentally determined force-dependent dissociation parameters for DNA-unzipping (19) and α_5_β_1_ integrin – RGD (FNIII_7–10_) interaction (21) were used in this study. We used the Bell-Evans energy landscape model with ssDNA elasticity correction to model DNA unzipping (19). (Figure 2a, b) In order to simulate the force-dependent dissociation of different DNA sequences, the dependencies of the unfolding energy barrier height 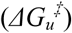and width 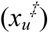on the length and CG content were extrapolated. (Figure 2c, see Supplementary Information for details) Our simulation produces excellent agreement with analytical solutions under both constant force and constant loading rate conditions. (Figure 2d, e). With this simulation tool, we are able to obtain bond rupture kinetics and statistics at high temporal resolution for systems of adhesion bonds subjected to an arbitrary mechanical history.

**Figure 2.**
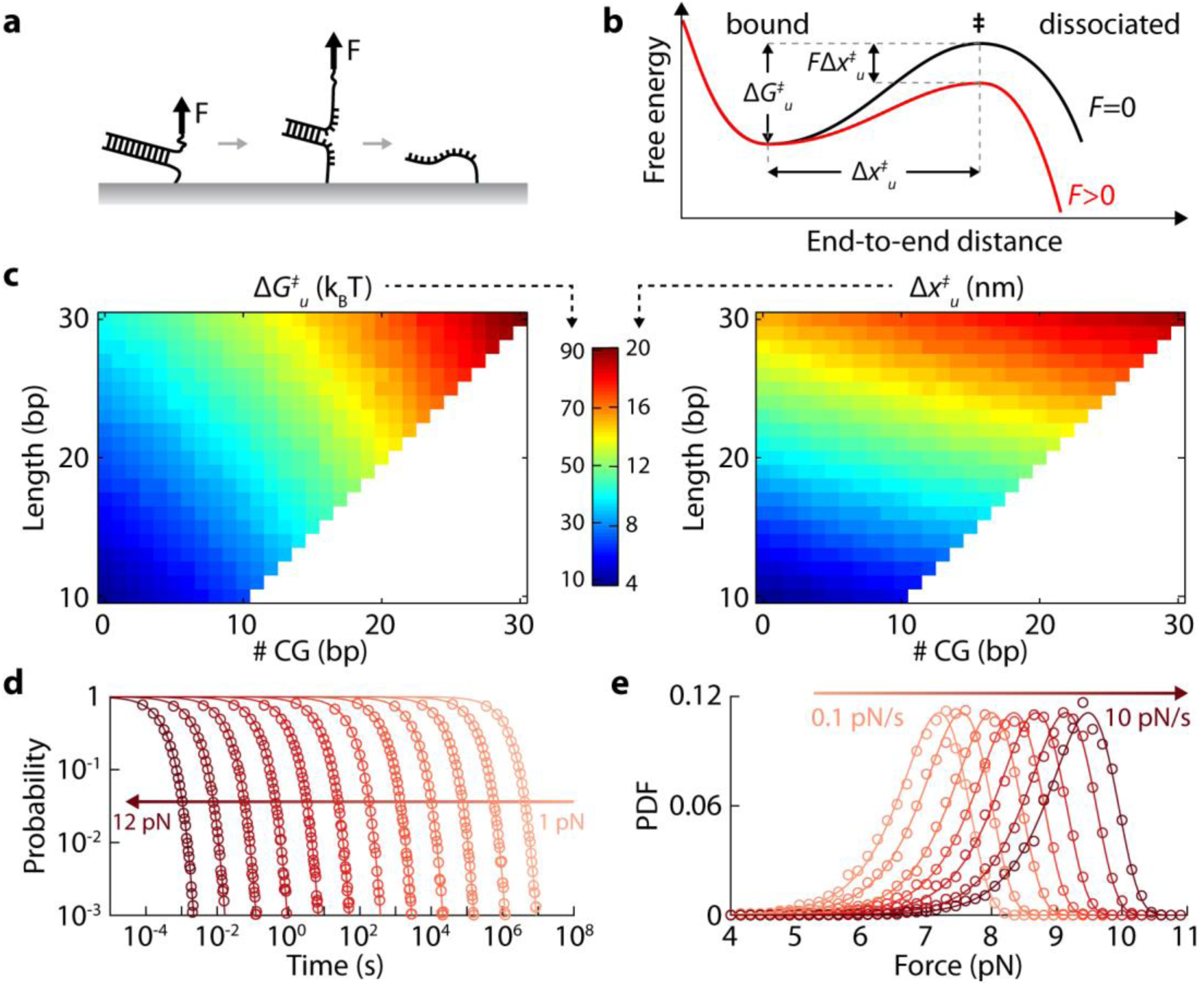
Monte Carlo simulation and validation of DNA unzipping kinetics. **(a)** Schematic of DNA unzipping, one of the core force sensing elements in TGT. **(b)** Schematic of the force effect on the energy landscape of DNA unzipping. **(c)** The DNA unzipping barrier height 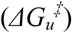and width 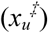as a function of DNA length (bp) and number of CG (bp). **(d)** Survival probability of intact DNA (L16/CG6) as a function of time over a range of constant forces (1 pN – 12 pN). **(e)** Probability density of DNA (L16/CG6) rupture force over a range of force loading rate (0.1 pN/s – 10 pN/s). Circles indicate results from simulation, solid lines indicate analytical solutions.

### TGT signal is modulated by receptor-ligand bond and is non-unique

The force-fluorescence response of the sensing element in a molecular force sensor is usually characterized separately by single-molecule force spectroscopy. It is assumed that the force-fluorescence response remains identical when the sensing element is linked to the target receptor. This is a valid assumption under biologically relevant time scales for reversible force sensors, as the sensing element is in a conformational dynamic equilibrium and is able to report instantaneous force via fluorescence. However, quantitative interpretation of the fluorescence signal generated from an irreversible force sensor is challenging because it is a cumulative signal that depends on several factors that cannot be easily teased apart. To illustrate this, we simulated the time-and force-dependent unzipping of DNA (L20/CG10), both in the presence and absence of a conjugated RGD peptides bound to α_5_β_1_ integrins. The unzipping of DNAs can be experimentally observed via fluorescence gains, while the rupture of the receptor-ligand bond keeps the DNA intact generating no fluorescence signals (Figure 3a, b) (17).

**Figure 3.**
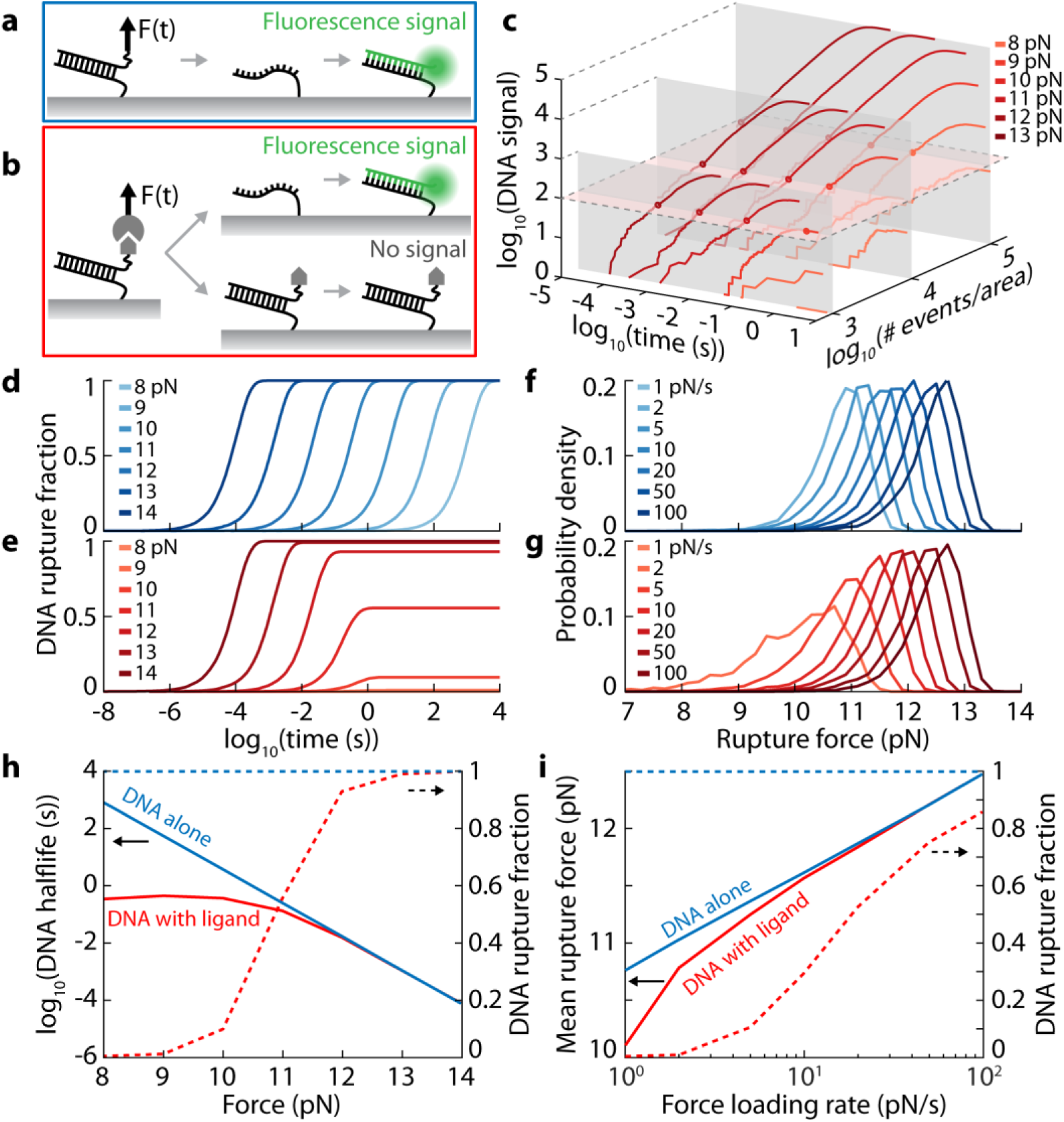
Simulation of DNA unzipping in the absence and presence of a receptor-ligand pair. **(a)** Fluorescence signal is generated whenever the top strand of the DNA is unzipped. **(b)** In the presence of a receptor-ligand pair, fluorescence signal is generated only in the outcome where DNA is unzipped. **(c)** Cumulative fluorescence signal per unit area as a function of force, time, and the surface density of adhesion events. Each dot on the same horizontal plane indicates a distinct combination of conditions that result in the same fluorescence intensity. **(d), (e)** The fraction of DNA rupture events as a function of time over a range of constant forces corresponding to scenario in (a) and (b), respectively. **(f), (g)** The probability density function of DNA rupture force over a range of loading rates, corresponding to scenarios in (a) and (b), respectively. **(h)** The apparent half-life and fraction of DNA rupture as a function of force, calculated from (d) and (e). **(i)** The mean DNA rupture force and fraction as a function of loading rate, calculated from (f) and (g). All DNA in the simulations are L20/CG10.

A constant force acting directly to unzip DNA (Figure 3a) will eventually unzip all the DNAs; as shown in Figure 3d, DNA rupture fraction converges to 1 for all forces. This is because force-dependent dissociation is irreversible and happens on the first-pass across the energy barrier. The barrier crossing rate increases with increasing force and is well described by previous models and experimental evidence (19). In this idealized system, if the surface density of TGT and the duration of force are known, the magnitude of force can theoretically be calculated.

In the case where forces are applied to stretch DNA together with the receptor/ligand pair (Figure 3b), the apparent DNA unzipping kinetics are modulated by the force-dependent rupture of the receptor-ligand bond as seen in Figure 3e. At higher forces (15 pN), DNA unzipping kinetics curve resembles that of DNA unzipping alone (Figure 3d), this is because the DNA lifetime at 15pN is significantly shorter than that of the receptor-ligand bond (Supp. Figure 1). As force decreases, the dissociation rate of DNA decreases and eventually drops below the dissociation rate of the receptor-ligand bond. Hence, the receptor-ligand bond rupture events contribute to the majority of all the rupture events while DNA unzipping makes up a small fraction (Figure 3e). Because of this, different force histories can produce the same level of fluorescence intensity during an imaging experiments. (Figure 3c) Therefore, quantitative interpretation of molecular forces based on fluorescence intensity alone is extremely challenging. In addition, the apparent kinetic half-life of DNA unzipping is significantly shortened in the presence of the receptor-ligand bond (Figure 3h). In an experiment where the only observable is fluorescence intensity change over time, the apparent lower DNA half-life would lead to an over-estimation of force. While the physical behaviour of the DNA unzipping is ideally independent of any attached receptor-ligand, the deviation is the result of receptor-ligand breaking event exhausting the total number of tethers under force. The effect is especially pronounced at lower molecular forces. Similarly, in constant loading rate scenarios, the apparent rupture force distribution in the presence of receptor-ligand is significantly skewed and shifted towards lower forces (Figure 3f, g). As force ramps up, an increasing number of receptor-ligand bonds break at lower forces, while the number of remaining DNA tethers available at higher forces decrease, causing a systematic suppression of DNA unzipping events at high force. In constant loading rate experiment, this would lead to an over-estimation of the rupture force; the effect is increasingly prominent at biologically relevant loading rates (22) below <10 pN/s (Figure 3i). Therefore, irreversible force sensors characterized in isolation can only be used to model fluorescence intensities of TGTs in experiments at high force and high loading rate regimes, where the force-dependent dissociation rate of the sensor is significant higher than that of the receptor-ligand pair.

Under both constant force and constant loading rate conditions, the fluorescence intensity resulting from DNA rupture is a function of force magnitude, duration, surface density, and the rupture characteristics of the receptor-ligand bond. Hence, a single snapshot of fluorescence image in cell experiment is insufficient to quantify force history. In addition, quantitative assessment TGT response in a real experimental system would require characterization of the force-dependent dissociation of the receptor-ligand bond through dynamic single-molecule force spectroscopy. This inevitably makes it difficult to use TGT as a modular component in various systems to study molecular adhesion.

### Force quantification via two force sensors in series

To address the challenges above, we investigated systems composed of two TGT elements in series (Figure 4a) and in parallel (Figure 4b). In the serial system, two different DNA sensor elements (DNA1, L16/CG6 and DNA2, L17/CG6) are serially connected to the receptor-ligand bond. When subjected to force, there are three possible outcomes (at t = ∞): (1) receptor-ligand bond ruptures; (2) DNA1 ruptures; or (3) DNA2 ruptures. The rupture events of DNA1 and DNA2 can be detected via fluorescently labeled (red and green) complementary ssDNA, where *R* and *G* are fractions of all the rupture events. We three systems under constant force (range in 0-15 pN): (1) without conjugate, (2) conjugated to RGD-α_5_β_1_ integrin, and (3) conjugated to DNA3 (L18/CG6 in the unzipping geometry). While the profiles of *R* and *G* are distinct across the three conditions (Figure 4c), all three showed identical *G/R* ratio, which is a one-to-one function of force (Figure 4e). When DNA1 and DNA2 are placed in series, the pair form an internal cross-reference that is undisturbed by the force-dependent rupture behaviour of the receptor-ligand bond. Therefore, the magnitude of force determined from *G/R* is completely independent of the receptor-ligand bond it probes, in contrast to the case of a single DNA. This makes it possible to apply serial DNA sensors calibrated by single-molecule force spectroscopy to any adhesion bonds knowing the calibration results will hold. Furthermore, the dimensionless *G/R* makes force quantification independent of surface density, which is another factor that readouts from individual TGTs cannot distinguish.

**Figure 4.**
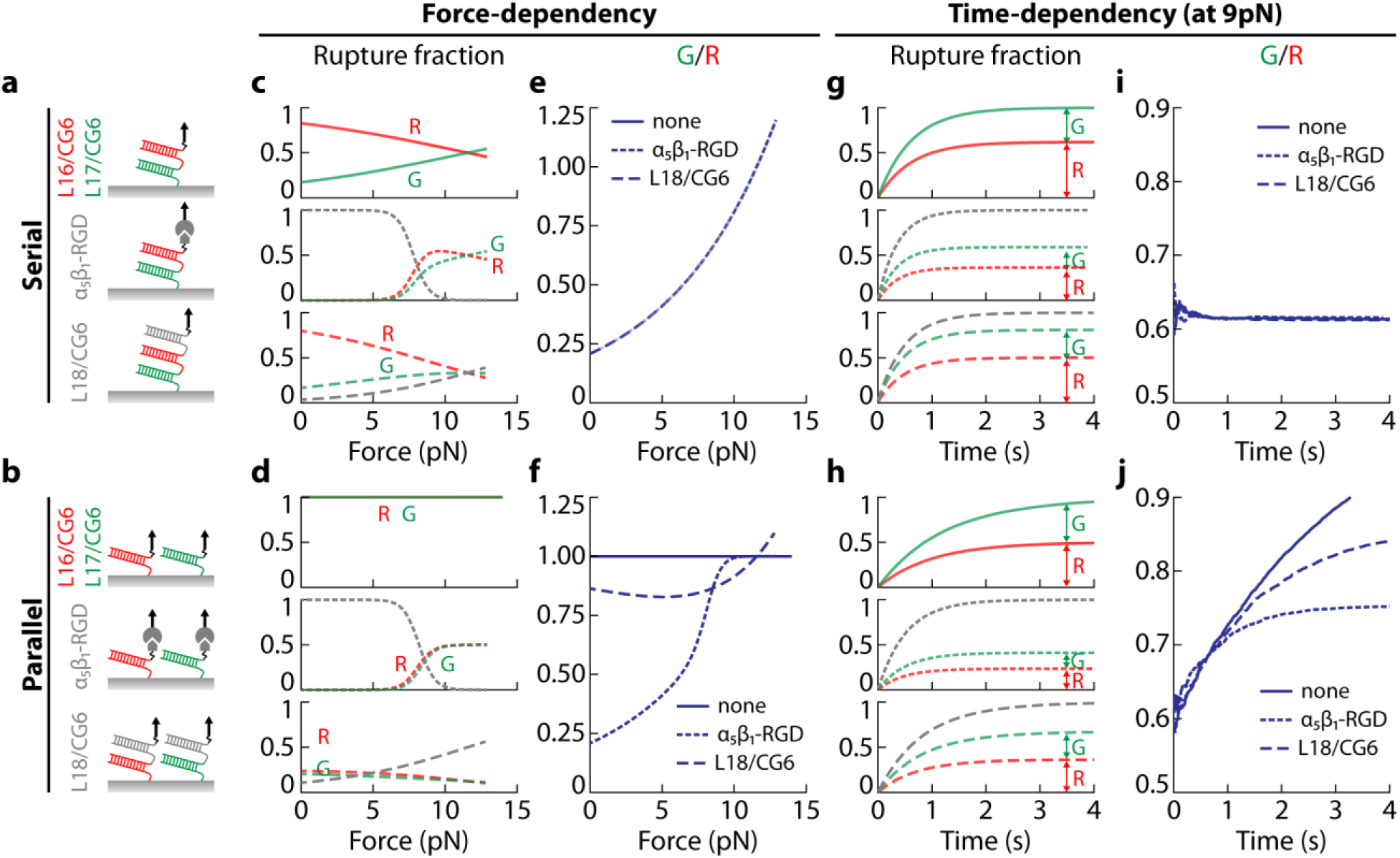
Force-and time-dependent signals from serial and parallel TGTs. **(a), (b)** Schematics of the serial and parallel configurations. The red and green DNAs (L16/CG6 and L17/CG6) together make up the sensing element and will produce red (*R*) and green (*G*) fluorescence signal when ruptured. The gray DNA (L18/CG6) and integrin-RGD are two distinct target adhesion bonds tested in this simulation. **(c), (d)** Rupture fraction of each component as a function of force at t = ∞, colours of lines match the components in (a) and (b). **(e), (f)** *G/R* as a function of force, with line style (solid, dashed, and dotted) indicate the three different cases, matching (c) and (d). **(g), (h)** Rupture fraction of each component (stacked) as a function of time at a constant force of 8 pN, colours match the components in (a) and (b). **(i), (j)** *G/R* as a function of force, with line style (solid, dashed, and dotted) indicate the three different cases, matching (g) and (h).

Next, we studied the time dependence of *G/R* under constant force conditions. The cumulative bond rupture events over time of each component are shown in Figure 4g. Even though individually, *R* and *G* are both strongly dependent on the connected receptor-ligand pair, *G/R* is time-invariant at any given force (Figure 4i). This implies that no matter when an observation is made or how long the constant force duration is, a single fluorescence snapshot will allow for force quantification with the only assumption being a constant force scenario. Therefore, *G/R* is a function of force and force alone, under constant force scenario. Because of this, if one assumes force is relatively constant in short time intervals, *ΔG/ΔR* can be used to report the magnitude of force in the time interval where *ΔG* and *ΔR* are incremental changes in fluorescence intensity. This would allow for force determination over time.

Lastly, we investigated the same pair of DNA sensors in a parallel configuration (Figure 4b). In this configuration, DNA1 and DNA2 are independently connected to the target receptor, which is often how multiplexing experiments were designed. We performed the same set of characterizations (Figure 4d, f, h,j) as we did for the serial configuration (Figure 4c, e, g, i). Our simulation shows that the force-dependent profile of *G/R* is dependent on the conjugated receptor-ligand pair (Figure 4f). Hence, this configuration suffers the same issues as the single DNA sensor system where the *G/R* is modulated by the targeted adhesion molecule modulation. Simulation of the time-dependent behaviour of the system (Figure 4h, j) shows that *G/R* changes over time in a conjugate-dependent manner. Therefore, the ratio of incremental fluorescence change *ΔG/ΔR* is also time-and conjugate-dependent. This does not come as a surprise as DNA1 and DNA2 rupture independently. There is no function *f(G,R,F)* in the parallel configuration that can make the result converge into one function as in the case of the serial configuration.

From the above results, two serially connected DNAs can be treated as a single force sensing element where force magnitude at a given time interval can be reported through the *ΔG/ΔR* ratio. The serial sensor design, for the first time, offers the possibility to use irreversible sensors to measure the magnitude of molecular force, which is not possible using a single irreversible sensor or through parallel multiplexing them.

### Probing force history and surface receptor density

Because the ratio of fluorescence signal change *ΔG/ΔR* is only dependent on force, a force history can be reconstructed from monitoring fluorescence changes over time. We simulated constant loading rate (1 pN/s), and oscillatory force (8.75 and 7.75 pN intervals) scenarios to demonstrate how sub-pN force resolution can be achieved using serial TGT designs. A total of 10^5^ serial TGT (L16/CG6 and L17/CG6) with RGD-α_5_β_1_ integrin were simulated in each scenario. Fluorescence signals *R* and *G* are recorded to mimic experimental observables, which are converted to *ΔG/ΔR* using 50 ms time intervals to reconstruct forces using the established calibration curve (Figure 4e) for this TGT pair. The results show that force histories reconstructed from *ΔG/ΔR* closely reproduce the actual force history (Figure 5a, b) and that sub-pN changes are clearly detectable.

**Figure 5.**
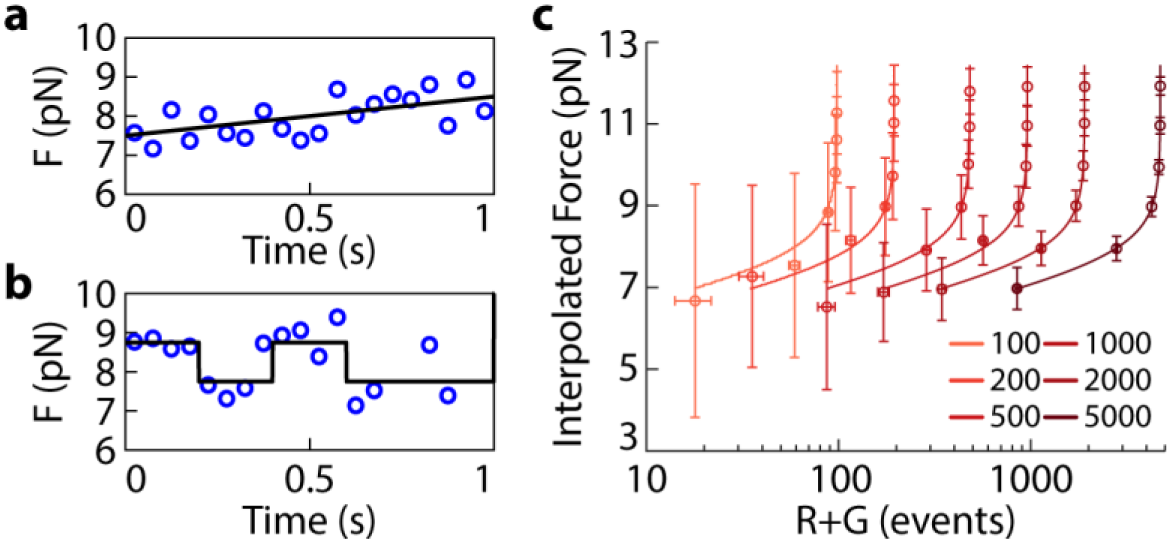
Force history and surface density reconstruction. **(a)** Reconstruction of force history from a constant loading rate simulation with the applied force starting at 7.5 pN and increasing at 1 pN/s. **(b)** Reconstruction of force history under an oscillatory force regime where the force alternated between 8.75 and 7.75 pN. **(c)** Illustration of the interplay between surface receptor density and the force applied to the system and the ability to determine both given a known number of (R+G) rupture events using the serial TGT design.

Next, we demonstrate that the receptor surface density can be calculated even though fluorescence signals from TGT rupture only report a fraction of adhesion events. One of the challenges in interpreting fluorescence signals from molecular force sensor is to tease apart contributions due to force and adhesion event density. This is particularly difficult in the case of a single TGT, as illustrated in Figure 3c, depending on the force history, only a fraction of TGT can rupture to produce fluorescence signal within a diffraction limited area. Serial TGT systems can bypasses this issue, as there is a singular solution that relates: (1) the magnitude of force, (2) the fluorescence intensity, and (3) the number of adhesion events within the same unit area. Given any force (or *ΔG/ΔR*) within a time interval, the total number of adhesion events in this time interval is proportional to *ΔG+ΔR*, where the proportionality constant dependent on the magnitude of the force only (Figure 5c). Hence, knowing *ΔG* and *ΔR* allows for determination of both the force magnitude and surface density of adhesion event.

### Rational design of serial force probes

There is a large DNA sequence space to explore for serial TGT designs. Since the behaviour of force-dependent DNA unzipping can be well characterized by the length and CG content of DNA, these are the only two parameters used to develop a serial TGT design strategy. We systematically evaluated every possible pairs of DNA with length ranging from 15 bp to 30 bp and CG content from 0% to 100%.

First, we looked at how length and CG content affect the force-dependent lifetime τ of a single DNA before the irreversible unzipping occurs (Supp. Figure 2a-c). The slope of *log(τ)* vs. *F* plot is dictated by the unzipping energy barrier width, while a vertical shift is dictated by the ratio of barrier height and width (Figure 2b). The energy barrier width is primarily controlled by the length of the DNA, while the barrier height is a function of both length and sequence. Hence, increasing CG content while keeping the DNA length constant shifts *log(τ)* higher; while changing length controls the tilt of *log(τ)*. Supp. Figure 2d-f compares the *log(τ) vs. F* curve of L20/CG10 DNA against all DNAs with length range 10-30 bp and CG content 0-100%. As expected, similar length and CG content brings the *log(τ) vs. F* curve closer to that of L20/CG10 DNA as measured by the mean square distance (MSD) between the curves (Supp.Figure 2e, f). For each pair of DNA with RGD-integrin, we defined a force quantification range (*FQR = F*_*max*_ *-F*_*min*_) where force quantification is feasible through fluoresce ratio (*ΔG/ΔR*). There are three factors that restrict the FQR for each DNA pair: (1) Insufficient fluorescence signal due to RGD-integrin bond breaking. At low forces, RGD-integrin bond dissociation dominates all the dissociation events, yielding a low fluorescence signal that sets the lower limit of the force detection range. We defined the lower limit as the force where DNA unzipping makes up less than 10% of all the dissociation events (Figure 6b). (2)The force sensitivity of the DNA pair. If the change of *ΔG/ΔR* as a function of force is insignificant, force cannot be accurately reconstructed. In addition, fluorescence signal from both DNA must be detectable. We defined the useful range for a DNA pair as the rupture fraction (*f = R/(R+G)*) of either DNA1 (*R*) or DNA2 (*G*) must change at least 1% per 1 pN change in force, i.e. *df/dF > 0.01* pN^-1^ (Figure 6c). (3) The upper limit of unzipping rate that can be predicted using energy landscape theory. This defines the upper limit where the force tilts the energy landscape far enough that there is no longer an energy barrier between the folded and unfolded structure. DNA unzipping lifetime cannot be predicted above this point and we decided use this as the upper limit of FQR (Figure 6b). Combining the above three constraints, the overall *F*_*min*_ and *F*_*max*_ defines the FQR (Figure 6c).

**Figure 6.**
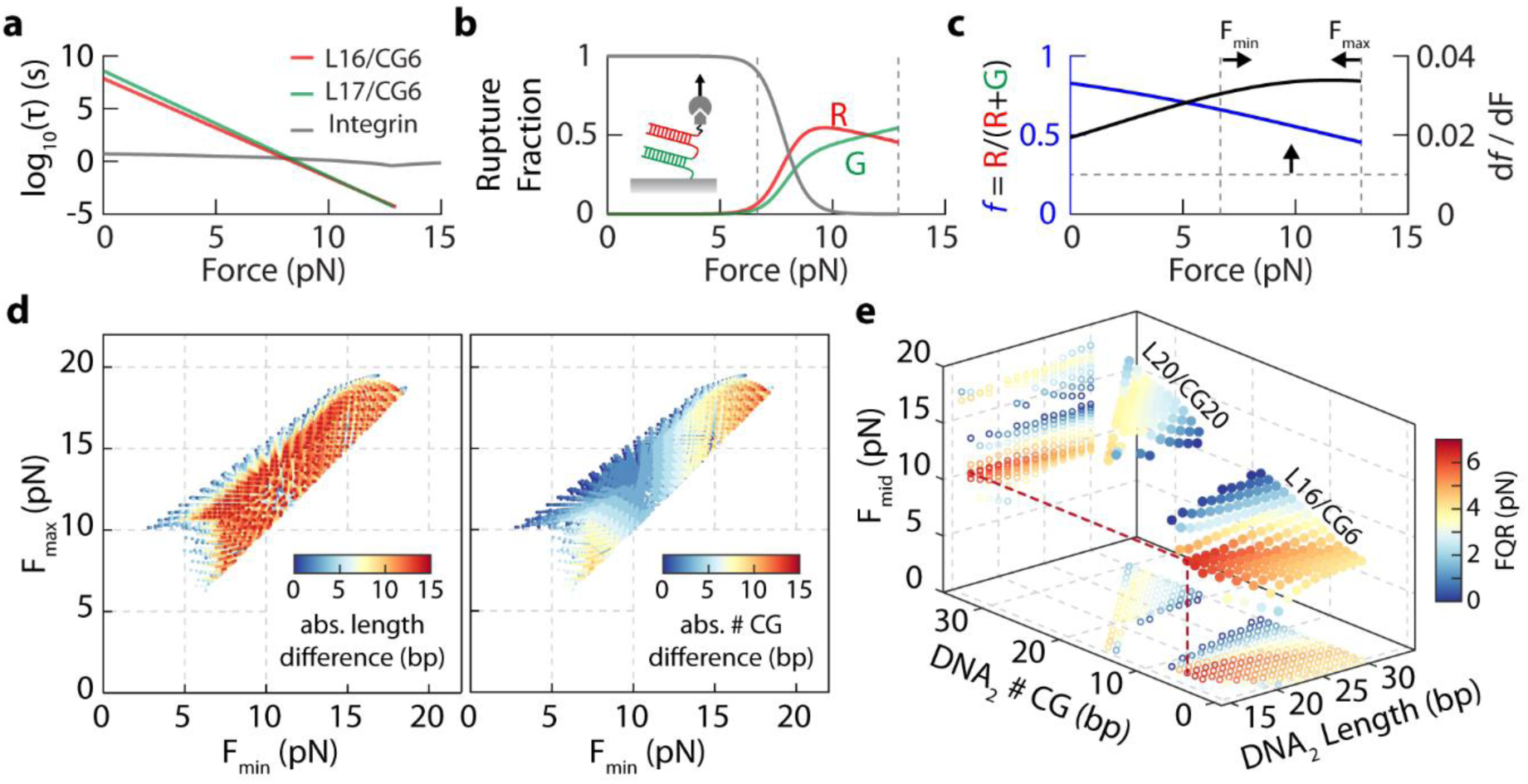
Systematic characterization of DNA pairs for serial TGTs. **(a)** Force-dependent dissociation lifetime of L16/CG6 (red), L17/CG6 (green), and RGD-integrin (gray). **(b)** Rupture fraction of each component (L16/CG6, L17/CG6, and RGD-integrin) in a serial TGT system at different forces. Gray dashed lines indicate the constraints from 10% DNA rupture events (left) and energy landscape prediction limit (right) **(c)** Fraction of DNA1 rupture among all DNA ruptures (*f=R/(R+G)*) and force sensitivity (*df/dF*) of serial TGT system as a function of force. The horizontal dashed line indicates the force sensitivity threshold for *df/dF* > 0.01/pN. The overall Fmin and Fmax are defined by the region of *df/dF* within the 3 constraints marked by the dashed lines. **(d)** Scatter plot of Fmax vs Fmin of serial TGT for every pair of DNA with length between 15 and 30 bp. The colour map indicates the length difference (left) and # of CG difference (right) of the DNA pair. **(e)** 3D scatter plot showing how Fmid and FQR depend on DNA2 sequence composition, for two DNA1 examples: L20/CG20, and L16/CG6.

To understand what factors controls *F*_*min*_ and *F*_*max*_, we looked at the differences in length and in number of CG bases between the two serially connected DNA (Figure 6d). Each dot in Figure 6d represents one pair of DNAs, with its location indicating its *F*_*min*_ and *F*_*max*_. The large spread shows the large tunability of the serial TGT system – there are pairs that can quantify in the 2-10 pN range on the low-end and 15-20 pN range on the high-end. The DNA pair with larger FQR tend to be ones with smaller length and CG differences (Figure 6d). This is because a few base-pairs of difference in length and CG is generally sufficient to produce *ΔG* and *ΔR* that respond distinctively at different forces, yet the sequence difference is not so large to ensure that both *ΔG* and *ΔR* are experimentally observable over a large force range. For illustration purposes, we picked two sequences for DNA_1_: L20/CG20 and L16/CG6 and systematically looked at how DNA_2_ sequences affect the FQR. In Figure 6e, we plotted the force at mid-range *F*_*mid*_*=(F*_*min*_*+F*_*max*_*)/2* as a function of the second pair’s length and number of CG, and colour-coded each dot by its FQR. Increasing CG while keeping length constant generally increases *F*_*mid*_ (Figure 6e), which is contributed to the up-shift of the log(τ) curve (Supp. Figure 2c) of DNA_2_. On the other hand, keeping the number of CGs in DNA2 constant while changing its length does little to Fmid, but affects the FQR significantly (Figure 6e). As the sequence composition of DNA_2_ approaches that of DNA_1_, FQR increases sharply. For L16/CG6, the sequence of DNA2 that gives rise to the highest FQR is F17/CG7; and L16/CG16 for L20/CG20.

## Discussion

Interpretation of fluorescence readout from irreversible force sensors such as the TGT requires great care. A single threshold force is likely insufficient to describe the force-dependent behaviour of molecular dissociation. For instance, in the unzipping geometry, L15/CG15 is “weaker” than L30/CG15 below ~12 pN, as shown by its lower unzipping lifetime τ (Supp. Figure 2a). However, at forces above ~12 pN, L15/CG15 becomes “stronger” than L30/CG15. The crossover point at ~12 pN for these two DNAs defines the force at which the two are equality strong with equal dissociation probability. Hence, the relative strength of different TGTs cannot be compared simply by a single threshold force assigned for each. Similarly, when a single TGT is tethered to a receptor-ligand bond, the crossover force at which the lifetime of the TGT and receptor-ligand equals may serve as a better definition of the characteristic force that is specific to the TGT/receptor-ligand pair. Furthermore, we showed here that the apparent behaviour of TGT is modulated by how receptor-ligand bonds respond to force. These make it difficult to (1) quantitatively interpret fluorescence images from TGT and (2) to quantitatively compare forces using the same TGT across different receptors. By placing two distinct TGTs in series, they experience the same force and forms an internally calibrated pair. Relative to each other, their rupture probability ratio is directly related to the force and is entirely independent of the receptor-ligand bond.

The serial sensor design scheme allows us to detect molecular forces using population statistics rather than individual sensors. This is the first time a design involving irreversible processes is able to provide an analog force readout, which has tremendous advantages in studying molecular forces. Because of the vast candidate choices and design possibilities utilizing molecular dissociations, there is significant advantage using serial sensor designs to achieve drastically different sensor dynamic ranges.

How does the serial TGT design compare to reversible force sensors? One of the challenges of reversible sensor when used alone is the inability to tease apart the surface density of force events and the magnitude of force, even with the assumption of homogeneous force per unit area (e.g. within a diffraction limited pixel). For instance, 10% sensors subjected to 10 pN force may produce the same fluorescence readout as 20% sensors subjected to 5 pN force. This can be tackled (23) using a design where one can detect simultaneously the force and know how many sensors are being pulled. By having two distinct sensors in one construct – one that responses to low force, hence any force application shows that it has been pulled and gives information about the surface density, while the other sensor provides more quantitative force information. This type of design usually utilizes quencher and fluorophore systems and may suffer from sensitivity issues especially if the force event surface density is low. In addition, such system may not be most suitable for fast dynamics cellular events such as those in rolling adhesion where the receptor is relatively sparse and the force application is brief (10s – 100s of ms (17, 24)). A very high sensitivity, yet quantitative solution is required. The TGT system is most suitable for such single-adhesion, high-speed adhesion events as illustrated in our previous work. Using this design, we hope to resolve this issue and provide quantitative information about the mean molecular force required for rolling adhesion. The application for the serial TGT is hence most suitable for such system.

## Conclusion

The involvement of mechanical processes in a wide array of biological processes makes their study and quantification an increasingly important in the life sciences. Many methods have been developed and tested in the literature, each with their advantages and drawbacks. Here, we have introduced a new design to surmount some of the common issues faced when using molecular force sensors and validated it using Monte Carlo simulations. We have shown it to be independent of the adhesion molecule being studied as well as the surface receptor density. This design allows for the force, as well as the force history, to be determined for a wide range of biologically relevant adhesion molecules.

## Acknowledgments

This work is supported by funding from the National Science and Engineering Research Council (NSERC) of Canada, and the Canada Foundation for Innovation (CFI).

## Author contribution

YM and IL designed the research, YM carried out the simulation, YM and IL analyzed the data, YM and IL drafted the manuscript.

